# Humanized *Mcl-1* mice enable accurate pre-clinical evaluation of MCL-1 inhibitors destined for clinical use

**DOI:** 10.1101/335430

**Authors:** Margs S. Brennan, Catherine Chang, Grant Dewson, Lin Tai, Guillaume Lessene, Andreas Strasser, Gemma L. Kelly, Marco J. Herold

**Affiliations:** The Walter and Eliza Hall Institute of Medical Research, Melbourne 3052, Australia.; Department of Medical Biology, University of Melbourne, Melbourne 3010, Australia.

**Keywords:** MCL-1, BH3-mimetics, apoptosis, lymphoma, pre-clinical models

## Abstract

MCL-1 is a pro-survival BCL-2 protein required for the sustained growth of many cancers. Recently a highly specific MCL-1-inhibitor, S63845, showing 6-fold higher affinity to human compared to mouse MCL-1 has been described. To accurately test efficacy and tolerability of this BH3 mimetic drug in pre-clinical cancer models, we developed a humanized *Mcl-1 (huMcl-1)* mouse in which MCL-1 was replaced with its human homologue. *HuMcl-1* mice are phenotypically indistinguishable from wild-type mice but are more sensitive to MCL-1 inhibition. Importantly, non-transformed cells and lymphomas from *huMcl-1;Eμ-Myc* mice are more sensitive to S63845 *in vitro* than their control counterparts. When *huMcl-1;Eμ-Myc* lymphoma cells are transplanted into *huMcl-1* mice, treatment with S63845 alone or alongside cyclophosphamide leads to long-term remission in ~60% or almost 100% of mice, respectively. These results demonstrate the potential of our huMCL-1 mouse model to test MCL-1 inhibitors, allowing precise predictions of efficacy and tolerability for clinical translation.

## INTRODUCTION

Apoptosis is a form of programmed cell death required for the removal of unwanted and potentially dangerous (e.g. infected) cells in multicellular organisms (Green and Llambi, 2015). Deregulation of the intrinsic apoptotic pathway can lead to accumulation of cells that should normally be deleted, including those with DNA lesions. The abnormal proliferation of damaged and mutated “undead” cells, can lead to tumour development. Cell death hinges on the balance between the pro-survival and pro-apoptotic members of the BCL-2 family of proteins. Defined by the presence of at least one of four BCL-2 homology (BH) domains, this family is further divided into the pro-survival members (BCL-2, BCL-X_L_, BCL-W, MCL-1 and BCL2A1/BFL-1), the BH3-only pro-apoptotic proteins (such as BIM and PUMA) and the multi BH domain pro-apoptotic effectors (BAK, BAX and BOK). Diverse stress stimuli cause increased expression of BH3-only family members to initiate apoptosis, either indirectly by binding to the pro-survival BCL-2 proteins, thereby unleashing the pro-apoptotic effectors BAK and BAX, or by binding BAK and BAX directly. Activation of BAK and BAX leads to their oligomerization on the outer mitochondrial membrane, perforating its surface, thereby releasing cytochrome-c and other apoptogenic factors. This induces a cascade of signalling events driven by the caspases that causes demolition of the cell.

Cancer cells can subvert the apoptotic machinery by either up-regulation of pro-survival proteins or down-regulation of pro-apoptotic proteins, thereby maximizing their growth potential (Hanahan and Weinberg, 2011). MCL-1 is important for the sustained growth of many cancers. The *MCL-1* locus is amplified in numerous human tumor types (Beroukhim et al., 2010) and functional studies in mouse models have showed that MCL-1 is essential for the sustained growth of many cancers, including multiple myeloma (Gong et al., 2017), T cell lymphomas (Grabow et al., 2014; Spinner et al., 2016), MYC- (Kelly et al., 2014) or BCR-ABL-driven (Koss et al., 2013) pre-B/ B cell lymphomas, acute myeloid leukaemia (Glaser et al., 2012) as well as some subtypes of solid tumours (Xiao et al., 2015; Zhang et al., 2011). Direct targeting of pro-survival BCL-2 family members by so-called ‘BH3 mimetics’ is a successful approach in cancer therapy. The BCL-2 inhibitor Venetoclax is highly efficacious for the treatment of relapsed/refractory Chronic Lymphocytic Leukemia (CLL) (Roberts et al., 2016; Stilgenbauer et al., 2016). A clinically relevant specific and potent inhibitor of MCL-1, called S63845 (Kotschy et al., 2016), has been shown to be efficacious as a single agent in several pre-clinical models of haematological malignancies and in combination with oncogenic kinase inhibitors in certain lung, skin as well as breast cancer derived cell lines and patient-derived xenograft models (Kotschy et al., 2016; Merino et al., 2017).

The clinical application of such an MCL-1 inhibitor has been of great concern given the important role MCL-1 plays in many normal tissues. MCL-1 is widely expressed (Kozopas et al., 1993), and essential for embryonic development with homozygous loss of *Mcl-1* in mice resulting in failure to implant at the blastocyst stage (Rinkenberger et al., 2000). Furthermore, conditional gene knockout studies have shown that MCL-1 plays a vital role in the survival of cardiomyocytes (Thomas et al., 2013; Wang et al., 2013), hematopoietic stem cells (Opferman et al., 2005), developing and mature lymphocytes (Dzhagalov et al., 2008; Opferman et al., 2003; Peperzak et al., 2013; Vikstrom et al., 2010) and in the maintenance of oocytes in the ovarian reserve (Omari et al., 2015). In spite of the many important roles MCL-1 plays, very little toxicity was observed when healthy mice were treated *in vivo* with the MCL-1 inhibitor S63845 at a dose that ablates mouse lymphoma cells (Kotschy et al., 2016). One caveat of these results is that S63845 has a ~6-fold higher affinity for the human protein in comparison to murine MCL-1 and therefore it is possible that the mouse models utilized were not sensitive enough to reveal all potential on-target toxicities. It was therefore pivotal to test more accurately the action of S63845 in a pre-clinical model in which we had humanized the *Mcl-1* locus *(huMcl-1)* by replacing the native coding region of the murine *Mcl-1* locus with the human MCL-1 coding sequence. These mice are healthy and fertile, and the intrinsic apoptotic pathway is intact in their cells, demonstrating that the human MCL-1 protein can functionally fully replace the murine protein. As predicted, cells from huMcl-1 mice were more sensitive to S63845 compared to wild-type cells, but not other cytotoxic agents that induce killing by the apoptotic pathway. In line with these results the maximum tolerated dose of S63845 was lower in the *huMcl-1* mice compared to their wild-type counterparts. However, it was still possible to cure *huMcl-1* mice of MYC-driven lymphomas expressing human MCL-1 by treatment with S63845, either alone or at lower doses together with cyclophosphamide, without causing overt damage to healthy tissues. These findings demonstrate the utility of the *huMcl-1* model to accurately determine the efficacy and tolerability of S63845 and potentially other MCL-1 inhibitory drugs in pre-clinical models of disease.

## RESULTS

### Generation of the humanised Mcl-1 mouse model

The humanized *Mcl-1* (*huMcl-1*) mouse model was generated by standard gene targeting in C57BL/6 embryonic stem (ES) cells. Expression of the human MCL-1 protein under the control of the murine *Mcl-1* promoter and regulatory regions was achieved by substitution of the exons and interspersing introns of the mouse *Mcl-1* locus with the human *MCL-1* exon/intron sequences, while maintaining the flanking 5’ and 3’ untranslated region of the genomic locus of the mouse (Fig. 1A). Homozygous loss of *Mcl-1* causes embryonic lethality prior to embryonic-day 4 in mice (Rinkenberger et al., 2000). Furthermore, changes in the 5’ untranslated region of the murine *Mcl-1* locus can lead to infertility in homozygous male *Mcl-1* floxed mice (Okamoto et al., 2014). Importantly, we found that the humanization of MCL-1 in mice had no impact on fertility or embryonic development, with the expected Mendelian ratios and sex distribution observed in intercrosses of *Mcl-1^hu/wt^* mice (Fig. 1B). Western blot analysis of thymocytes from *huMcl-1* and wild-type mice showed efficient expression of the human MCL-1 protein, as evidenced by the predicted size difference between the mouse and human MCL-1 proteins (human: 38 kD, mouse: 35 kD, Fig. 1C). A cohort of *huMcl-1* mice was aged for >600 days without showing any obvious signs of defects.

**Figure 1.**
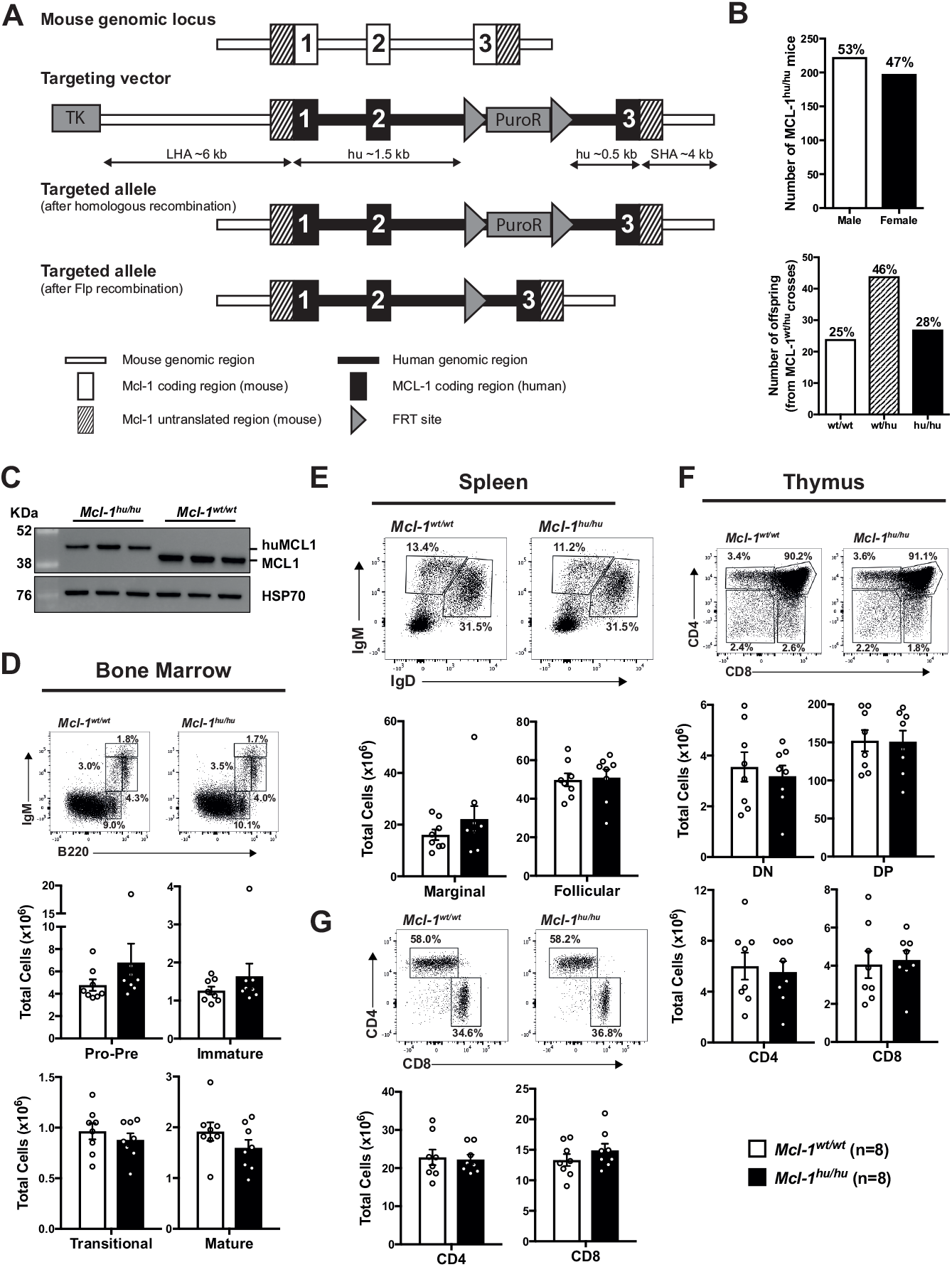
Humanized *Mcl-1* mice show no defects under steady state conditions. (**A**) Schematic representation of the murine *Mcl-1* gene locus, humanised *Mcl-1* targeting vector and the correctly targeted alleles, before and after Flpe recombination. (**B**) Offspring sex frequency of *Mcl-1^hu/hu^* mice and Mendelian ratios from *Mcl-1^wt/hu^* heterozygous matings. (**C**) Thymocytes were isolated from *Mcl-1^hu/hu^* and *Mcl-1^wt/wt^* mice. MCL-1 protein (mouse and human) expression was detected by Western blotting. Probing for HSP70 served as a loading control. (**D-G**) Single cell suspensions were prepared from spleen, thymus and bone marrow of *Mcl-1^hu/hu^* and *Mcl-1^wt/wt^* mice and cell subsets were determined by immunostaining and FACS analysis. Data are presented as mean ± SEM, significance determined by students t-test. (**D**) Representative FACS plot of B cell development in the bone marrow (top) and total cell numbers per femur (bottom panels). Pro-B/pre-B (B220^lo^IgM^−^), immature B (B220^lo^IgM^mid^), transitional B (B220^lo-hi^IgM^hi^) and mature (B220^hi^IgM^mid^) B cells. (**E**) Representative FACS plot of peripheral B cells in the spleen (top) and total cell number (bottom). B cells defined as follicular (IgM^+^IgD^+^) and marginal zone (IgM^+^IgD^lo^) B cells. (**F**) Representative FACS plot depicting T cell development in the thymus (top) and total cell numbers calculated for each population (bottom panels). Immature double negative thymocytes (DN, CD4^−^CD8^−^), double positive thymocytes (DP, CD4^+^CD8^+^) and the mature CD4 and CD8 single positive populations. (**G**) Representative FACS plot of peripheral T cell distribution (top) in the spleen as well as total cell numbers of CD4 and CD8 single positive T cell populations (bottom panel).

### Humanised Mcl-1 mice have normal hematopoietic cell subset distribution

MCL-1 is crucial for the survival of many developing and mature cell subsets of the hematopoietic system in mice (Dzhagalov et al., 2008; Huntington et al., 2007; Opferman et al., 2005; Opferman et al., 2003; Peperzak et al., 2013; Vikstrom et al., 2010). FACS analysis of the bone marrow revealed no differences in the frequencies and numbers of pro-B/pre-B (B220^lo^IgM^−^), immature B (B220^lo^IgM^+^), transitional B (B220^mid^IgM^hi^) or mature B (B220^hi^IgM^lo^) cell subsets between *huMcl-1* mice and wild-type controls (representative FACS plots shown; Fig. 1D). There were also no abnormalities in IgM^+^ or IgD^+^ peripheral B lymphocytes in the *huMcl-1* mice (Fig. 1E). The frequencies and numbers of the different stages of thymocyte development, as determined by staining for CD4 and CD8, were also comparable between *huMcl-1* mice and wild-type controls (representative FACS plot shown; Fig. 1F). Moreover, *huMcl-1* mice had normal frequencies and numbers of mature CD4^+^ and CD8^+^ T lymphocytes in the spleen (Fig. 1G), with normal distributions of naïve (CD62L^+^CD44^−^), effector memory (CD62L^−^CD44^+^) and central memory (CD62L^+^CD44^+^) CD4^+^ T cells (Fig. S1A). Finally, the *huMcl-1* mice had normal frequencies and numbers of macrophage/monocyte (MAC-1^+^GR-1^lo^) and neutrophil populations (MAC-1+GR-1^hi^) in the bone marrow and spleen (Fig. S1B).

### Human MCL-1 protein can functionally replace mouse MCL-1 within the apoptotic machinery

Since it is possible that other BCL-2 proteins (pro-apoptotic or pro-survival) are differentially expressed to compensate for the expression of huMCL-1 instead of the mouse MCL-1 protein *in vivo*, we determined the levels of these proteins by Western blotting. Thymocytes and splenocytes from *huMcl-1* mice showed no difference in the protein levels of the pro-apoptotic BIM and PUMA or pro-survival BCL-2, BCL-XL or A1 (Representative Western blot Fig. 2A, multiple blots quantified in Fig. 2B). The affinity for MCL-1 antibodies to the mouse versus human protein is not known, thus we could not compare the relative protein expression by Western blot. To test whether the huMCL-1 protein expressed in mouse cells is able to interact with the endogenous mouse pro-apoptotic BCL-2 relatives, we performed co-immunoprecipitation assays on thymocyte extracts. This revealed that the huMCL-1 protein expressed in mouse cells is capable of binding to mouse BIM and BAK in a similar manner as mouse MCL-1 (Fig. 2C). To assess the functionality of the huMCL-1 protein, we investigated the response of cells from the *huMcl-1* mice to a diverse range of cytotoxic stimuli *in vitro*. Importantly, no abnormalities were observed in the survival of thymocytes or B cells from the *huMcl-1* mice (Fig. 2D). As predicted, thymocytes and B cells from the *huMcl-1* mice were more sensitive to the MCL-1 inhibitor S63845 compared to those from wild-type mice (Fig. 2E). While the increased sensitivity to S63845 became only obvious at higher (200 nM, 1 μM) doses in thymocytes, more striking differences were observed in B cells after 6 hours of treatment with low (40 nM) or high (1 μM) doses of the MCL-1 inhibitor (Fig. 2E, 24h Fig. S2A).

**Figure 2.**
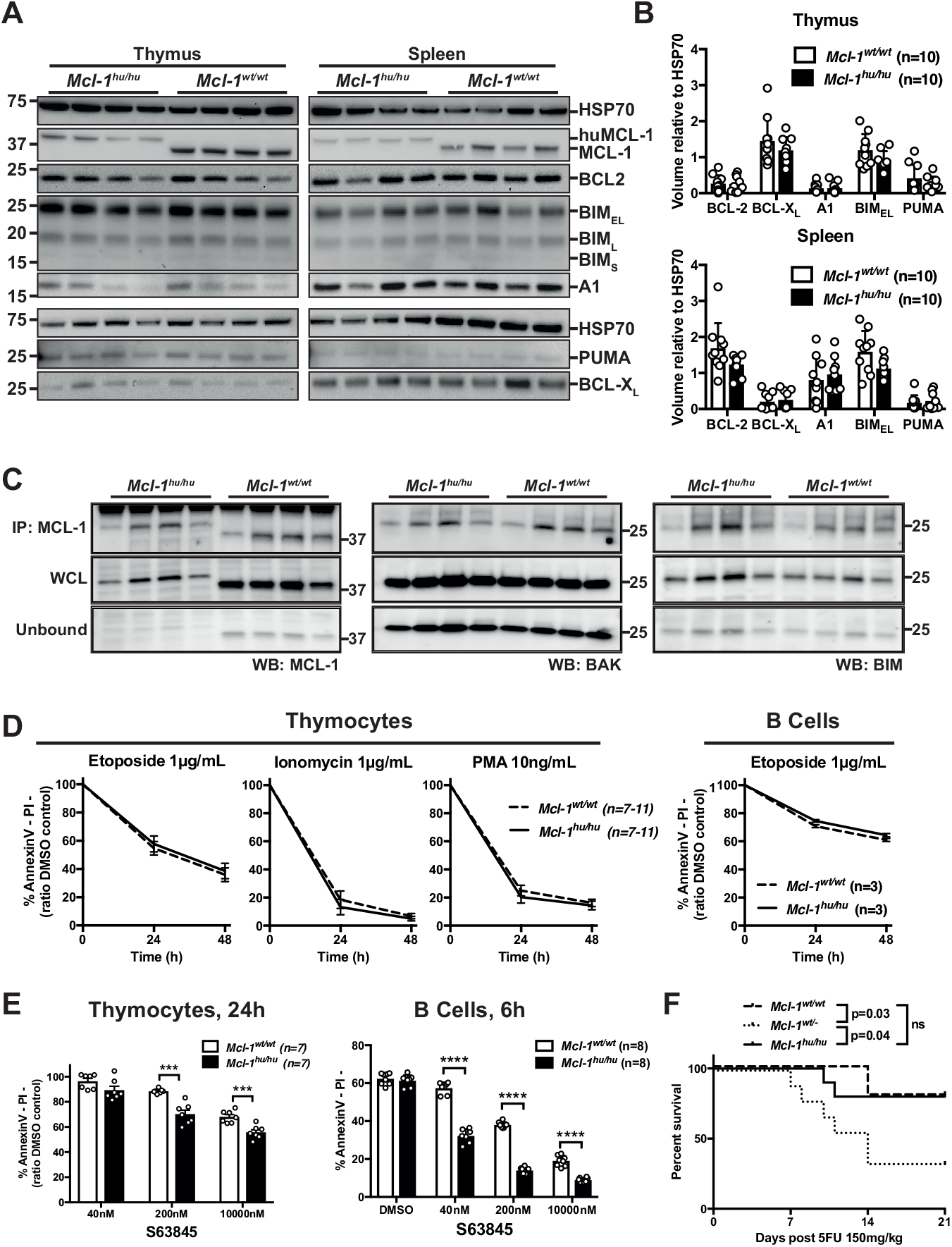
Apoptosis pathway remains intact in humanized *Mcl-1* mice. (**A**) Thymocytes and splenocytes were isolated from *Mcl-1^hu/hu^* and *Mcl-1^wt/wt^* mice. Expression of MCL-1, BCL-2, BCL-XL, A1, BIM and PUMA proteins was detected by Western blotting. Probing for HSP70 served as a loading control. Representative blot is shown. (**B**) Quantification of expression levels of BCL-2 family proteins in thymocytes and splenocytes from *huMcl-1* (n=9) and wild-type (n=10) mice determined by Western blotting. Chemiluminescent signals from each protein were normalized to the HSP70 loading control signal and plotted as arbitrary units; significance calculated by multiple t-tests. (**C**) Lysates were prepared from thymocytes from *huMcl-1* and wild-type mice and immunoblotted for MCL-1 (left panel), BAX (middle panel) and BIM (right panel) after immunoprecipitating with an MCL-1 (binds both the mouse and human MCL-1 proteins) specific antibody. Pulled-down fraction shown in top panels with whole cell lysate (WCL) and unbound fractions shown below. (**D**) Thymocytes and splenic B cells were isolated from *huMcl-1* and wild-type mice and cultured with a variety of cytotoxic stimuli as indicated. Cell viability at 24 and 48 h was assessed by AnnexinV/PI staining and FACS analysis. (**E**) Thymocytes and splenic B cells were isolated from *huMcl-1* and wild-type mice and treated with the MCL-1 inhibitor S63845 at the indicted concentrations. Cell viability at 24 h (and also 6 h for B cells) was assessed by AnnexinV/PI staining and FACS analysis. Data are presented as mean ± sem; significance calculated by multiple t-tests at each time point, *=p>0.05, **=p>0.01, ***=p>0.001, ***=p>0.0001, ns=not significant. (**F**) Mice of the indicated genotypes were treated with a single dose of 150 mg/kg body weight Fluorouracil (5FU) and survival monitored. Kaplan-Meier curve; statistical analysis was performed by log rank (Mantel-Cox) test, *=p>0.05.

Next, we wanted to test the ability of huMCL-1 protein expressed in mouse cells to function within a whole animal. MCL-1 is required for the maintenance of hematopoietic stem/progenitor cells (Opferman et al., 2005) and for emergency hematopoiesis following myeloablative chemotherapy (Delbridge et al., 2015). The latter was demonstrated when *Mcl-1^wt/-^* mice were found to be compromised in their recovery from 5-fluorouracil (5-FU) treatment. We treated *huMcl-1, Mcl-1^wt/-^* (as a control) and wild-type mice with a single dose of 5-FU (150 mg/kg body weight) and monitored for hematopoietic recovery over a period of 21 days. As reported (Delbridge et al., 2015), most *Mcl-1^wt/-^* mice did not recover from 5-FU treatment, whereas *huMcl-1* mice recovered as well as wild-type controls (Fig 2F). Collectively, these findings demonstrate that huMCL-1 is functional and the apoptotic machinery is intact in cells from the *huMcl-1* mice, both *in vitro* and *in vivo*. Furthermore, as predicted, these cells show increased sensitivity to the MCL-1 inhibitor S63845.

### Determining the maximum tolerated dose of S63845 in humanized Mcl-1 mice

Given the binding affinity of S63845 is ~6-fold higher for human MCL-1 protein compared to mouse MCL-1, we determined the maximum tolerated dose (MTD) of S63845 in the *huMcl-1* mice. Treatment of *huMcl-1* mice on 5 consecutive days intravenously (i.v.) with doses ranging from 5 to 25 mg/kg body weight established the MTD of S63845 at 12.5 mg/kg (Fig 3A). At 15 mg/kg S63845, 1 out of 4 mice did not survive while none of the mice could tolerate the drug at 25 mg/kg. The tolerability of the drug was considerably higher in wild-type mice, which all survived the 25 mg/kg dosing schedule (Fig.3A) and have previously been shown to have an MTD of 40 mg/kg (Kotschy et al., 2016). This demonstrates that the *huMcl-1* mice are more sensitive to the MCL-1 inhibitor S63845.

**Figure 3.**
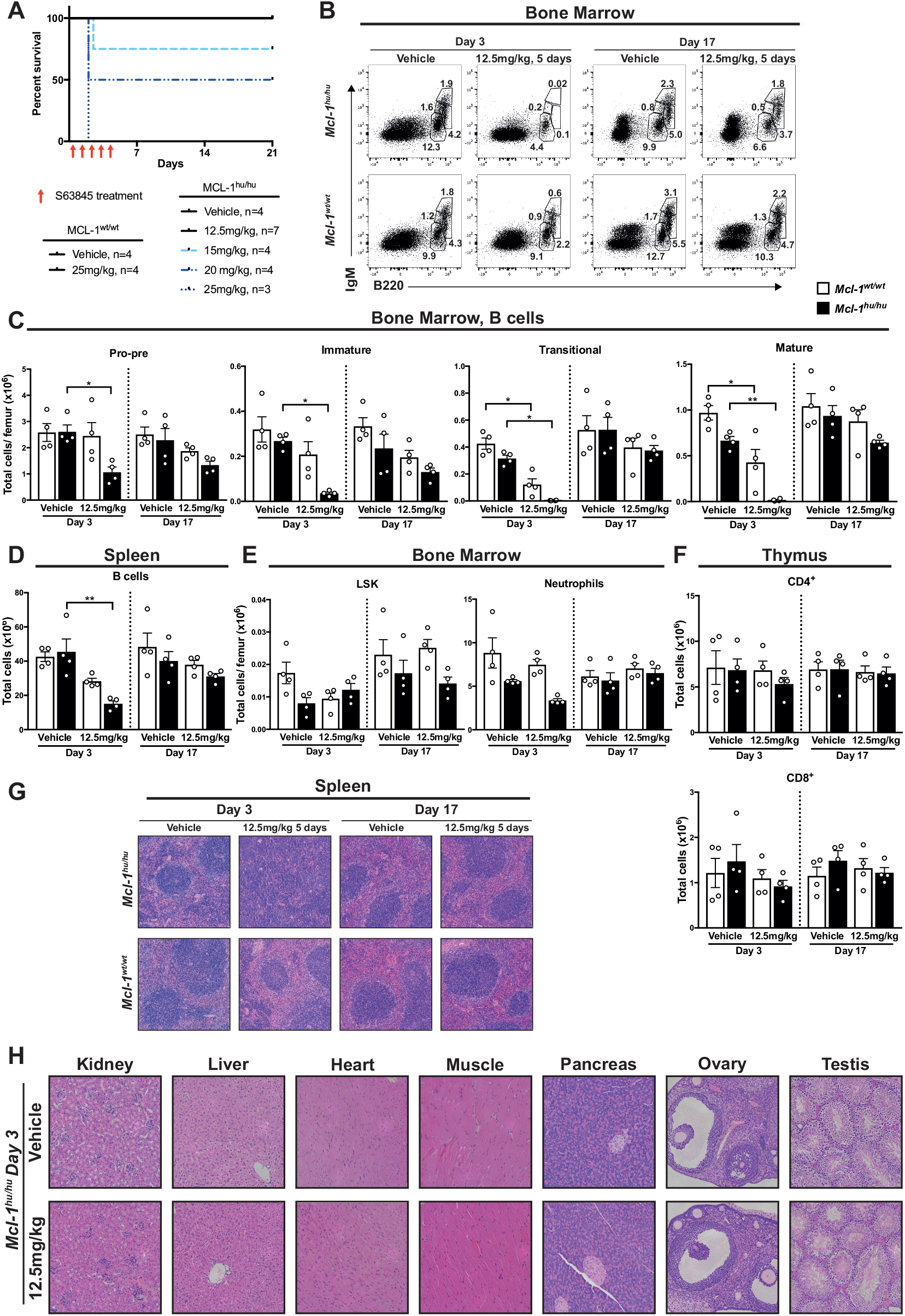
S63845 exerts no enduring toxicity in the *huMcl-1* mouse model at the MTD. (**A**) Kaplan-Meier survival curves of *huMcl-1* or wild-type mice treated for 5 consecutive days with vehicle or S63845 with increasing doses, as indicated. (**B-H**) *huMcl-1* and wild-type mice were treated with 5 consecutive doses of 12.5 mg/kg S63845 or vehicle. 3 or 17 days post-treatment organs were harvested for analysis. (**B-F**) Single cell suspensions were prepared from spleen, thymus and bone marrow of *Mcl-1^hu/hu^* and *Mcl-1^wt/wt^* mice and cell subsets were determined by immunostaining and FACS analysis. Data are presented as mean ± SEM, significance determined by students t-test. (**B**) Representative FACS plot of B cells in the bone marrow and total cell numbers per femur (**C**). B cells defined as pro-B/pre-B (B220^lo^IgM^−^), immature B (B220^lo^IgM^mid^), transitional B (B220^lo-hi^IgM^hi^) and mature B cells (B220^hi^IgM^mid^). (**D**)Total cell numbers of B cells in the spleen (B220^+^) (**E**) LSK cells (Lineage^−^SCA-1^+^c-KIT^+^) and macrophages/monocytes (MAC-1^+^GR-1^lo^) in the bone marrow (**F**) and mature T cells (CD4^+^ or CD8^+^) in the thymus. (**G**) Representative H&E stained section of spleens from *huMcl-1* or wild-type mice that had been treated with 12.5 mg/kg S63845 or vehicle and were harvested at either 3 or 17 days after treatment. (**H**) Representative H&E stained sections of various organs as indicated from *huMcl-1* mice treated with 12.5 mg/kg S63845 or vehicle 3 days after treatment had ceased.

### S63845 exerts no enduring toxicity in the huMcl-1 mice at the MTD

There have been many reports on the importance of MCL-1 in several vital tissues (Dzhagalov et al., 2008; Omari et al., 2015; Opferman et al., 2005; Opferman et al., 2003; Peperzak et al., 2013; Thomas et al., 2013; Vikstrom et al., 2010; Wang et al., 2013) raising the question whether an MCL-1 inhibitor would find its therapeutic use in the clinic. Previously it was shown that S63845 exerts very little toxic effects in mice expressing mouse MCL-1 (Kotschy et al., 2016), but given the higher affinity of this drug for human MCL-1 we revisited this question by treating *huMcl-1* (and wild-type controls) mice for 5 consecutive days with either vehicle or 12.5 mg/kg S63845 and testing for acute impact and recovery 3 or 17 days post treatment, respectively. In the *huMcl-1* mice, the pro-B/pre-B (B220^lo^IgM^−^), immature B (B220^lo^IgM^+^), transitional B (B220^mid^IgM^hi^) and mature B (B220^hi^IgM^lo^) B cells in the bone marrow were significantly reduced at 3 days but had mostly recovered at 17 days post treatment (representative FACS plots Fig. 3B; total cell numbers Fig. 3C). Similar observations were made for B cells in the blood (Fig. S3A) and spleen (Fig. 3D). Despite the reduction in splenic B cells, an increase in cellularity was noted in the spleens of all *huMcl-1* mice treated with S63845 3 days after the treatment, but these values returned to normal at 17 days post treatment (Fig. S3B). FACS analysis revealed that the increased splenic cellularity was due to an increase in EryA (Lineage^−^Ter119^hi^CD71^+^FSC-A^hi^) and EryB (Lineage-Ter119^hi^CD71^+^FSC-A^lo^) erythrocyte progenitor populations, with the more mature EryC (Lineage^−^Ter119^hi^CD71^−^FSC-A^to^) remaining stable throughout the experiment (representative FACS plots Fig. S3C; total cell numbers Fig. S3D). Moreover, the RBC counts in the peripheral blood showed a modest reduction at day 3 post treatment, but again these numbers were completely recovered at day 17 (Fig.S3E). No significant changes were observed in the LSK (Lineage^−^SCA-1^+^c-KIT^+^, Fig. 3E), or neutrophil (MAC-1^+^GR-1^hi^) populations in the bone marrow (Fig. 3E) or spleen (Fig. S3F) of the S63845 treated *huMcl-1* mice. Moreover, there were no changes in the numbers of mature CD4^+^ and CD8^+^ T cells (Fig 3F), immature double negative (DN1-4, Lineage^−^TCRb^−^ CD4^−^CD8^−^) and double positive (DP, CD4^+^CD8^+^) thymocytes (Fig. S3G) or mature T cells (TCRß^+^) in the spleen of S63845 treated *huMcl-1* mice (Fig S3G). Histological analysis revealed an increased size and loss of normal architecture of the spleen, which complements the FACS data (Fig. 3G). Histological analysis of major organs revealed no damage in response to S63845 (Fig. 3F). These data indicate that S63845 treatment can be tolerated in *huMcl-1* mice, with only a transient reduction of certain hematopoietic cell subsets.

### Eμ-Myc cell lines expressing huMcl-1 are more sensitive to S63845

In order to test whether the sensitivity towards MCL-1 inhibition with S63845 could be increased not only in healthy but also malignant cells, we crossed the *huMcl-1* mice with the *Eμ-Myc* transgenic animals. While the latency and tumour phenotype did not differ between mouse or human MCL-1 expressing *Eμ-Myc* mice (Fig. 4A-B), the *in vitro* sensitivity of cell lines generated from sick mice increased by 6-fold in the *Eμ-Myc;huMcl-1* lymphoma cells towards MCL-1 inhibition with S63845 compared to control *Eμ-Myc* lymphoma cells expressing mouse MCL-1 protein (25nM or 160nM respectively, Fig. 4C-D).

**Figure 4.**
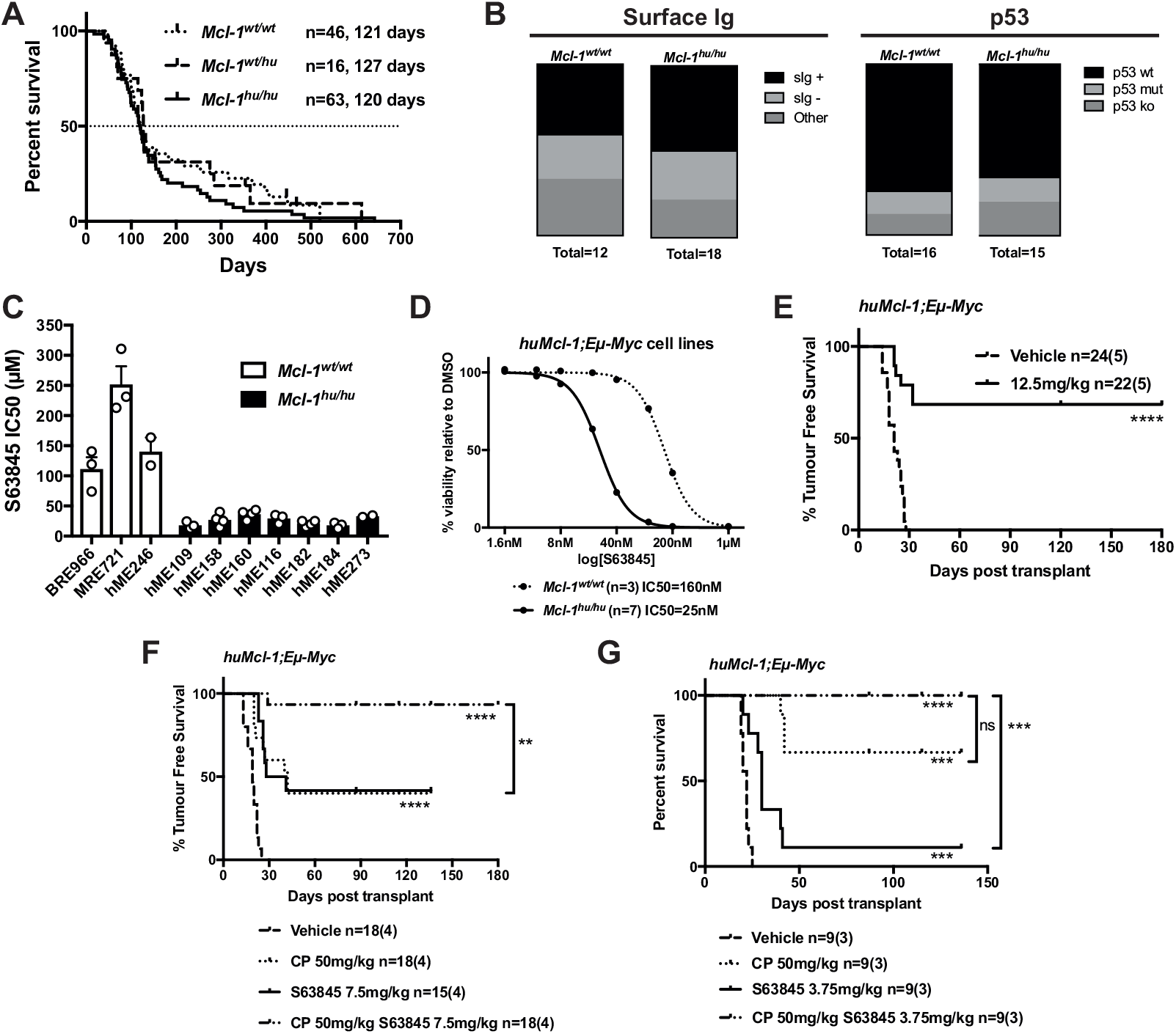
*HuMcl-1;Eμ-Myc* lymphoma cells are more sensitive to S63845 treatment compared to control *Eu-Myc* lymphoma cells. (**A**) *Eμ-Myc* transgenic mice that were*Mcl-1^wt/wt^, Mcl-1^wt/hu^* or *Mcl-1^hu/hu^* were aged and monitored for tumor-free survival. Kaplan-Meyer survival curve shown with median latency for each genotype indicated. (**B**) Primary wild-type or *huMcl-1 Eμ-Myc* lymphoma cells were isolated and immunostained for surface Ig (sIg) expression (IgD and/or IgM) and analyzed by FACS. p53 and p19 expression were detected by Western blotting. Proportions of p53 wild-type (p53 low-negative, p19 low-negative) p53 knock-out (p53 negative, p19 high) and p53 mutant (p53 high, p19 high) are shown. (**C**) Viability of *Eμ-Myc* lymphoma cells that are wild-type for *Mcl-1* (n=3, white bars) or homozygous for the *huMcl-1* allele (n=7, black bars). Cell viability was determined by AnnexinV/PI staining and FACS analysis. Data points are IC50 values calculated from a range of concentrations (4 nM-1 μM) analyzed in triplicate. (**D**) Cell viability data generated and shown in (**C**) were pooled to compare the overall IC50 of control *Eμ-Myc* and *huMcl-1;Eμ-Myc* lymphoma cell lines. Combined IC50 curves for each genotype are shown, with IC50 values depicted in the legend. (**E-G**) Survival curves of *huMcl-1;Ly5.1* mice transplanted with *huMcl-1;Eμ-Myc* lymphomas and treated with vehicle, S63845 alone (doses as indicated) for 5 consecutive days (**E**), or S63845 in combination with cyclophosphamide (CP, **F-G**). n= number of recipient mice with number of cell lines in brackets. Significance relative to vehicle shown in graph; significance relative to combination treatment shown to the right. Calculated by Mantel-Cox test, *=p>0.05, **=p>0.01, ***=p>0.001, ****=p>0.0001, ns=not significant.

### Regression of huMcl-1;Eμ-Myc lymphomas in vivo with S36845, either alone or in combination with cyclophosphamide

To test the sensitivity of *Eμ-Myc* lymphomas expressing human MCL-1 to S63845 *in vivo*, we transplanted *huMcl-1;Eμ-Myc* lymphoma cell lines into *huMcl-1;Ly5.1* recipient mice (i.e. both lymphoma and normal cells expressed huMCL-1). After 3 days, mice were treated for 5 consecutive days with vehicle or 12.5 mg/kg S63845 and monitored for signs of sickness. 60% of mice were cured at this dose (Fig. 4E, survival curves of individual cell lines; Fig. S4A). Next, we aimed to improve tumour free survival in mice transplanted with *huMcl-1;Eμ-Myc* lymphoma cell lines. To this end, *huMcl-1;Eμ-Myc* lymphoma cell lines were transplanted into *huMcl-1;Ly5.1* recipient mice and 2 days later treated with vehicle or a low dose of cyclophosphamide (CP, 50 mg/kg). Three days later mice were treated for 5 consecutive days with a low dose of S63845 (7.5 mg/kg). Treatment with 50mg/kg CP or 7.5mg/kg S63845 by themselves resulted in ~50% or ~25% tumor free survival, respectively. Excitingly, only one mouse became sick with lymphoma in the combination treatment increasing the efficacy of S63845 thereby increasing the tumor free survival to almost 100% (Fig. 4F, individual cell lines: Fig. S4B). For cell lines that did previously regress at 12.5mg/kg bodyweight S63845 (hME160, hME184, hME273; Fig. S4A), cancer free survival was also achieved with as little as 3.75mg/kg bodyweight S63845 in combination with CP (Fig 4G). These findings, in a highly relevant mouse model in which both lymphoma and normal cells express huMCL-1, clearly demonstrate that there is a therapeutic window for S63845, and that its efficacy can be enhanced by combining treatment with cyclophosphamide.

## DISCUSSION

Here we describe a novel humanized *Mcl-1* mouse model, in which human MCL-1 protein is expressed under the control of the mouse *Mcl-1* regulatory regions. Normal embryonic development and fertility of these mice is noteworthy given that even small changes to the *Mcl-1* locus or loss of only one allele bear consequences in adult mice (Okamoto et al., 2014), with homozygous loss of *Mcl-1* being embryonic lethal very early in embryonic development (Rinkenberger et al., 2000). Moreover, the intrinsic apoptotic pathway remains intact in cells from the *huMcl-1* mice with no compensatory changes in the levels of other BCL-2 family members detected (Fig. 2). This is critical because changes in expression of the BCL-2 family member proteins can compensate for the loss of another. For example, tumor cells can acquire resistance to inhibitors of BCL-2, BCL-XL and BCL-W (e.g. ABT-737) by up-regulating MCL-1 (Lin et al., 2007). Since the huMcl-1 mice are normal and their cells have an unimpaired apoptotic machinery (confirmed both *in vitro* and *in vivo)* they represent an ideal model to predict tolerability and efficacy of MCL-1 inhibitors.

MCL-1 is regarded as an exciting therapeutic target due to its importance for the sustained growth of many cancers, and S63845 has been developed as a highly selective and potent inhibitor. A so far unique feature for BH3 mimetics is that S63845 binds more tightly to the human MCL-1 protein compared to the mouse MCL-1 protein and one could imagine that any newly generated compounds targeting MCL-1 would likely bind in the same region, and therefore, will likely have similar properties. Since previous *in vivo* experiments have all been conducted with mouse MCL-1 as the target, it was difficult to determine whether the results are truly reflective of the therapeutic potential of S63845 and perhaps more importantly, the safety. Our studies using the *huMcl-1* mouse model support the notion that a therapeutic window for MCL-1 inhibitors might be established, with a transient loss of B cells being the most significant side effect observed at doses of S63845 that could halt lymphoma growth in a substantial fraction of mice. These data may be viewed conflicting with previous data showing the importance of MCL-1 in many essential normal cell types, such as cardiomyocytes. However, we believe that the lack of significant side effects is due to the transient action and hence inhibition of MCL-1 by the drug in contrast to the irreversible loss of MCL-1 elicited by genetic deletion. Additionally, it is not yet known whether S63845 is available in all organs throughout the mouse after its administration, which may further explain the differences seen between deletion *vs* inhibition of MCL-1. Further pharmacokinetic studies must be performed to address this question.

While S63845 appears to be relatively well tolerated as previously shown in mice expressing mouse MCL-1, we have shown that S63845’s tighter binding to human MCL-1 is relevant *in vivo* with the MTD of S63845 in *huMcl-1* mice being ~3 times lower than that of wild-type mice. This suggests that the *huMcl-1* mouse model will be useful for future pre-clinical work using S63845 (and other MCL-1 inhibitors) for determining a therapeutic window for treatment of cancers and possibly other diseases in which MCL-1 is important. As a proof of principal, we generated *huMcl-1;Eμ-Myc* lymphomas and showed that they are more sensitive to S63845, both *in vitro* and *in vivo*, than *Eμ-Myc* lymphomas expressing mouse MCL-1. The observation that *huMcl-1* mice could tolerate doses of S63845 that could prevent growth of *huMcl-1;Eμ-Myc* lymphomas suggests that a therapeutic window of MCL-1 inhibitors may be established in the clinic. While there are already phase I clinical trials being conducted for MCL-1 inhibitors, it is important to generate relevant pre-clinical data in different cancer models using our *huMcl-1* mice to help make rational decisions on which malignancies will benefit the most in these early stages of clinical development, and to determine which other anti-cancer agents can cooperate with MCL-1 inhibitors in tumor cell killing and still be tolerable as a combination therapy. Of note, we are currently breeding the *huMcl-1* alleles onto the NOD/SCID/common-gamma chain knockout (NSG) background, which will allow the testing of MCL-1 on primary human cancer cells (xenografts) with human MCL-1 expression in the healthy host cells. These highly relevant pre-clinical models will help guide the safe and appropriate use of MCL-1 inhibitors in cancer patients.

## ACKNOWLEDGMENTS

We thank the Herold and Strasser labs, Andrew W. Roberts and David C. S. Huang for advice in the preparation of the manuscript, Giovanni Siciliano, Krystal Hughes, Dan Fayle, Hannah Johnson and Cassandra D’Alessandro for technical assistance for *in vivo* experiments and animal husbandry. Our work is supported by the Australian National Health and Medical Research Council (Project Grant 1145728 to MJH 1143105 to MJH and AS, 1086291 to GLK; Program Grant 1016701 to AS and Fellowship 1020363 to AS), the Leukemia and Lymphoma Society of America (LLS SCOR 7001-13 to AS and MJH), the Cancer Council of Victoria (1086157 and 1147328 to GLK, 1052309 to AS and Venture Grant MJH and AS), a VCA fellowship (MCRF17028 to GLK), a Leukaemia Foundation Grant in Aid (to AS and GLK) and a Postgraduate Award (to MSB), sponsored research funding from Servier, a bequest from the Estate of Antony Redstone (to AS and GLK) as well as by operational infrastructure grants through the Australian Government Independent Research Institute Infrastructure Support Scheme (9000220) and the Victorian State Government Operational Infrastructure Support Program.

## AUTHOR CONTRIBUTIONS

The experiments were conceived and designed by MSB, AS, GLK and MJH. Experiments were performed by MSB, CC, GD, LT and GLK. GL contributed expertise and provided reagents. The paper was written by MSB, AS, GLK and MJH with help from the other authors.

## STAR METHODS

### Generation of the humanized Mcl-1 mice

Mice bearing the *huMcl-1* allele were generated by Taconic Biosciences GmbH, Cologne, Germany. The targeting vector (designed to insert the human *MCL-1* coding sequence under the control of the endogenous *Mcl-1* promoter, Fig. 1A) was transfected into the TaconicArtemis C57BL/6NTac embryonic stem (ES) cell line. Homologous recombinant clones were isolated using positive (PuroR) and negative (Thymidine kinase - TK) selection. Correctly targeted ES cells were injected into BALB/c blastocysts that were then transferred into pseudo-pregnant NMRI females. Chimeric offspring were selected by coat color for further breeding. The *PuroR* sequence was flanked by FRT sites to allow subsequent removal of the selection cassette by crossing chimeric mice with the C57BL/6-Tg(*CAG-Flpe)2 Arte* transgenic mice carrying the FLP recombinase (FLPe), giving rise to the final *huMcl-1* allele. Genotyping was performed by PCR to confirm cassette removal, loss of the FLPe transgene, and presence of the wild-type or humanized *Mcl-1* alleles (For PCR primers see Supplementary Table 1).

### Co-immunoprecipitation and Western blotting

Thymocytes were isolated from *Mcl-1^wt/wt^* and *Mcl-1^hu/hu^* mice and prepared in lysis buffer (Supplementary Table S2) supplemented with protease inhibitors cOmplete and pepstatin A (Roche and Sigma). Cell lysates were collected and pre-cleared with sepharose beads for 1 h at 4°C with constant agitation. Immunoprecipitation was performed using 2.5 μg monoclonal antibody against MCL-1 (Supplementary Table S2) incubated overnight at 4°C, followed by incubation with protein G sepharose beads for 1 h at 4°C. Immunoprecipitated proteins were eluted by boiling in SDS-PAGE sample buffer for 5 min and analyzed by Western blotting (antibodies listed in Supplementary Table S2). For all other Western blots, cell lysates were prepared in RIPA buffer (Supplementary Table S3) supplemented with complete protease inhibitor (Roche). Protein concentration was determined by Bradford assay using the Protein Assay Dye Reagent Concentrate (Bio-Rad, Hercules, CA, USA). Samples of 15 mg protein were prepared in Laemmli buffer (Supplementary Table S3), boiled for 5 min and size fractionated by gel electrophoresis on NuPAGE 10% Bis-Tris 1.5 mm gels (Life Technologies) in MES buffer and then transferred onto nitrocellulose membranes (Life Technologies) using the iBlot membrane transfer system. Antibody dilution and blocking were performed in 5% skim milk, 0.1% Tween 20 in PBS. For antibodies refer to Supplementary Table S3, in house antibodies (Lang et al., 2014; Okamoto et al., 2014). Luminata Forte Western HRP substrate (Millipore, Billerica, MA, USA) was used for developing and membranes were imaged and analyzed using the ChemiDoc XRS^+^ machine with ImageLab software (Bio-Rad).

### Immunostaining andflow cytometry

Thymus, spleen and bone marrow were harvested and single cell suspensions prepared in PBS (Gibco), 5 mM EDTA (Merck), supplemented with 5% fetal bovine serum (FBS, Sigma-Aldrich) for staining. Monoclonal antibodies (Supplementary Table 4) were obtained from eBioscience, BioLegend or generated at the Walter and Eliza Hall Institute (WEHI) Antibody Facility. Streptavidin-PE (BioLegend) was used to detect biotinylated antibodies. Propidium iodide (PI, 1 μg/mL) was used to exclude dead cells. For steady state analysis (Fig. 1, 1S), Whole organ cell counts were determined by the CASY counter (Schärfe System GmbH)(Fig. 1, S1) or by mixing a known concentration of APC Calibrite beads (Becton Dickinson) with each sample (Fig. 3, 3S). Data were collected using LSR II, LSRFortessa or LSRFortessa X-20 analyzers and examined using FlowJo 10 (Becton Dickson).

### Tissue culture and cell viability assays

*Eμ-Myc* lymphoma cell lines were maintained as previously described (Kelly et al., 2014). For all experiments, viability was determined by resuspending cells in Annexin V binding buffer (0.1 M Hepes (pH 7.4), 1.4 M NaCl, 25 mM CaCl_2_) containing PI (1 μg/mL) and FITC- or Alexa Fluor 647-conjugated Annexin V (generated in house). Spleen and thymi were harvested and single cell suspensions prepared. To isolate B cells, splenocytes were pelleted and resuspended in 1 mL red cell lysis buffer (156 mM ammonium chloride (BDH), 11.9mM sodium bicarbonate (Merck), EDTA (Sigma)) for 5 min to deplete erythrocytes. Cells were then stained with biotinylated antibodies against CD4, CD8, MAC-1 and GR-1 (Supplementary Table S4) to enrich for B cells by MagniSort Streptavidin Negative Selection Beads (Thermo Fisher), as per the manufacturer’s protocol. Isolated B cells and thymocytes were seeded at 5 × 10^4^ cells/well, in triplicate per condition, in 96 well flat-bottomed plates with either S63845 (8, 200, 1000 nM, gift from Servier), dexamethasone (1 nM), etoposide (1 □g/mL), PMA (10 ng/mL) or ionomycin (1 μg/mL) (Sigma) for the indicated times. The sensitivity of *Eμ-Myc* lymphoma cell lines to S63845 was determined by seeding 5 × 10^4^ lymphoma cells into 96 well flat-bottomed plates in FMA medium with 5 concentrations of S63845 (1:5 dilutions starting from 1 μM, gift from Servier), in triplicate and incubated at 37°C and 10% CO_2_ for 24 h. The IC_50_ values were determined using nonlinear regression algorithms in Prism (GraphPad).

### Animals and in vivo drug treatments

The care and use of mice for experimental purposes were carried out in accordance with the requirements set out by the Walter and Eliza Hall Institute (WEHI) Animal Ethics Committee. *HuMcl-1* mice are described above; *Eμ-Myc* (Adams et al., 1985) and *Mcl-1^wt/-^* (Vikstrom et al., 2010) mice have been described previously. All mice are kept on a C57BL/6-Ly5.2 background. The *huMcl-1* allele was bred onto a C57BL/6-Ly5.1 background for use as recipient animals for all transplant and toxicity experiments. Single cell suspensions of 1 × 10^5^ *Eμ-Myc* lymphoma cells in PBS were injected into 8-12 week old *huMCL-1;Ly5.1* recipient mice by intravenous (i.v.) tail vein injection. Recipient mice were sex matched to transplanted tumors. Working solutions of cyclophosphamide (CP) and 5-FU (Sigma) were prepared in PBS. CP was administered by intraperitoneal (i.p) injection, 5-FU by i.v. tail vein injection. S63845 (Servier) was formulated extemporaneously and protected from light in 2% Vitamin E/TPGS (Sigma) in NaCl 0.9% (w/v) and delivered by i.v. tail vein injection at indicated doses and schedule. Mice were monitored for sickness and killed when deemed unwell in order to generate survival curves by experienced mouse technicians (blinded to the treatments and genotypes of the mice).

### Statistical analysis

Prism software (GraphPad) was used to generate survival curves and perform all statistical testing of data. All data is presented as mean±s.e.m unless otherwise stated. P values > 0.05 were considered significant.

## REFERENCES

Adams, J.M., Harris, A.W., Pinkert, C.A., Corcoran, L.M., Alexander, W.S., Cory, S., Palmiter, R.D., and Brinster, R.L. (1985). The *c-myc* oncogene driven by immunoglobulin enhancers induces lymphoid malignancy in transgenic mice. Nature 318, 533–538.

Beroukhim, R., Mermel, C., Porter, D., Wei, G., Raychaudhuri, S., Donovan, J., Barretina, J., Boehm, J., Dobson, J., Urashima, M., et al. (2010). The landscape of somatic copy-number alteration across human cancers. Nature 463, 899–905.

Delbridge, A.R., Opferman, J.T., Grabow, S., and Strasser, A. (2015). Antagonism between MCL-1 and PUMA governs stem/progenitor cell survival during hematopoietic recovery from stress. Blood 125, 3273–3280.

Dzhagalov, I., Dunkle, A., and He, Y.W. (2008). The anti-apoptotic bcl-2 family member mcl-1 promotes T lymphocyte survival at multiple stages. J Immunol 181, 521–528.

Glaser, S., Lee, E.F., Trounson, E., Bouillet, P., Wei, A., Fairlie, W.D., Izon, D.J., Zuber, J., Rappaport, A.R., Herold, M.J., et al. (2012). Anti-apoptotic Mcl-1 is essential for the development and sustained growth of acute myeloid leukemia. Genes Dev 26, 120–125.

Gong, J.N., Khong, T., Segal, D., Yao, Y., Riffkin, C.D., Garnier, J.M., Khaw, S.L., Lessene, G., Spencer, A., Herold, M.J., et al. (2017). Hierarchy for targeting pro-survival BCL2 family proteins in multiple myeloma: pivotal role of MCL1. Blood, Epub ahead of print.

Grabow, S., Delbridge, A.R., Valente, L.J., and Strasser, A. (2014). MCL-1 but not BCL-XL is critical for the development and sustained expansion of thymic lymphoma in p53-deficient mice. Blood 124, 3939–3946.

Green, D.R., and Llambi, F. (2015). Cell Death Signaling. Cold Spring Harb Perspect Biol 7.

Hanahan, D., and Weinberg, R.A. (2011). Hallmarks of cancer: the next generation. Cell 144, 646–674.

Huntington, N.D., Puthalakath, H., Gunn, P., Naik, E., Michalak, E.M., Smyth, M.J., Tabarias, H., Degli-Esposti, M.A., Dewson, G., Willis, S.N., et al. (2007). Interleukin 15-mediated survival of natural killer cells is determined by interactions among Bim, Noxa and Mcl-1. Nat Immunol 8, 856–863.

Kelly, G.L., Grabow, S., Glaser, S.P., Fitzsimmons, L., Aubrey, B.J., Okamoto, T., Valente, L.J., Robati, M., Tai, L., Fairlie, W.D., et al. (2014). Targeting of MCL-1 kills MYC-driven mouse and human lymphomas even when they bear mutations in p53. Genes Dev 28, 58–70.

Koss, B., Morrison, J., Perciavalle, R.M., Singh, H., Rehg, J.E., Williams, R.T., and Opferman, J.T. (2013). Requirement for antiapoptotic MCL-1 in the survival of BCR-ABL B-lineage acute lymphoblastic leukemia. Blood 122, 1587–1598.

Kotschy, A., Szlavik, Z., Murray, J., Davidson, J., Maragno, A.L., Le Toumelin-Braizat, G., Chanrion, M., Kelly, G.L., Gong, J.N., Moujalled, D.M., et al. (2016). The MCL1 inhibitor S63845 is tolerable and effective in diverse cancer models. Nature 538, 477–482.

Kozopas, K.M., Yang, T., Buchan, H.L., Zhou, P., and Craig, R.W. (1993). *MCL1*, a gene expressed in programmed myeloid cell differentiation, has sequence similarity to *bcl-2*. Proceedings of the National Academy of Sciences of the United States of America 90, 3516–3520.

Lang, M.J., Brennan, M.S., O’Reilly, L.A., Ottina, E., Czabotar, P.E., Whitlock, E., Fairlie, W.D., Tai, L., Strasser, A., and Herold, M.J. (2014). Characterisation of a novel A1-specific monoclonal antibody. Cell death & disease 5, e1553.

Lin, X., Morgan-Lappe, S., Huang, X., Li, L., Zakula, D.M., Vernetti, L.A., Fesik, S.W., and Shen, Y. (2007). ‘Seed’ analysis of off-target siRNAs reveals an essential role of Mcl-1 in resistance to the small-molecule Bcl-2/Bcl-X(L) inhibitor ABT-737. Oncogene 26, 3972–3979.

Merino, D., Whittle, J.R., Vaillant, F., Serrano, A., Gong, J.N., Giner, G., Maragno, A.L., Chanrion, M., Schneider, E., Pal, B., et al. (2017). Synergistic action of the MCL-1 inhibitor S63845 with current therapies in preclinical models of triple-negative and HER2-amplified breast cancer. Science translational medicine 9.

Okamoto, T., Coultas, L., Metcalf, D., van Delft, M.F., Glaser, S.P., Takiguchi, M., Strasser, A., Bouillet, P., Adams, J.M., and Huang, D.C. (2014). Enhanced stability of Mcl1, a prosurvival Bcl2 relative, blunts stress-induced apoptosis, causes male sterility, and promotes tumorigenesis. Proceedings of the National Academy of Sciences of the United States of America 111, 261–266.

Omari, S., Waters, M., Naranian, T., Kim, K., Perumalsamy, A.L., Chi, M., Greenblatt, E., Moley, K.H., Opferman, J.T., and Jurisicova, A. (2015). Mcl-1 is a key regulator of the ovarian reserve. Cell Death Dis 6, e1755.

Opferman, J., Iwasaki, H., Ong, C.C., Suh, H., Mizuno, S., Akashi, K., and Korsmeyer, S.J. (2005). Obligate role of anti-apoptotic MCL-1 in the survival of hematopoietic stem cells. Science 307, 1101–1104.

Opferman, J.T., Letai, A., Beard, C., Sorcinelli, M.D., Ong, C.C., and Korsmeyer, S.J. (2003). Development and maintenance of B and T lymphocytes requires antiapoptotic MCL-1. Nature 426, 671–676.

Peperzak, V., Vikstrom, I., Walker, J., Glaser, S.P., LePage, M., Coquery, C.M., Erickson, L.D., Fairfax, K., Mackay, F., Strasser, A., et al. (2013). Mcl-1 is essential for the survival of plasma cells. Nat Immunol 14, 290–297.

Rinkenberger, J.L., Horning, S., Klocke, B., Roth, K., and Korsmeyer, S.J. (2000). Mcl-1 deficiency results in peri-implantation embryonic lethality. Genes Dev 14, 23–27.

Roberts, A.W., Davids, M.S., Pagel, J.M., Kahl, B.S., Puvvada, S.D., Gerecitano, J.F., Kipps, T.J., Anderson, M.A., Brown, J.R., Gressick, L., et al. (2016). Targeting BCL2 with Venetoclax in Relapsed Chronic Lymphocytic Leukemia. N Engl J Med 374, 311–322.

Spinner, S., Crispatzu, G., Yi, J.H., Munkhbaatar, E., Mayer, P., Hockendorf, U., Muller, N., Li, Z., Schader, T., Bendz, H., et al. (2016). Re-activation of mitochondrial apoptosis inhibits T-cell lymphoma survival and treatment resistance. Leukemia 30, 1520–1530.

Stilgenbauer, S., Eichhorst, B., Schetelig, J., Coutre, S., Seymour, J.F., Munir, T., Puvvada, S.D., Wendtner, C.M., Roberts, A.W., Jurczak, W., et al. (2016). Venetoclax in relapsed or refractory chronic lymphocytic leukaemia with 17p deletion: a multicentre, open-label, phase 2 study. The lancet oncology 17, 768–778.

Thomas, R.L., Roberts, D.J., Kubli, D.A., Lee, Y., Quinsay, M.N., Owens, J.B., Fischer, K.M., Sussman, M.A., Miyamoto, S., and Gustafsson, A.B. (2013). Loss of MCL-1 leads to impaired autophagy and rapid development of heart failure. Genes Dev 27, 1365–1377.

Vikstrom, I., Carotta, S., Luethje, K., Peperzak, V., Jost, P.J., Glaser, S., Busslinger, M., Bouillet, P., Strasser, A., Nutt, S.L., et al. (2010). Mcl-1 is essential for germinal center formation and B cell memory. Science 330, 1095–1099.

Wang, X., Bathina, M., Lynch, J., Koss, B., Calabrese, C., Frase, S., Schuetz, J.D., Rehg, J.E., and Opferman, J.T. (2013). Deletion of MCL-1 causes lethal cardiac failure and mitochondrial dysfunction. Genes Dev 27, 1351–1364.

Xiao, Y., Nimmer, P., Sheppard, G.S., Bruncko, M., Hessler, P., Lu, X., Roberts-Rapp, L., Pappano, W.N., Elmore, S.W., Souers, A.J., et al. (2015). MCL-1 Is a Key Determinant of Breast Cancer Cell Survival: Validation of MCL-1 Dependency Utilizing a Highly Selective Small Molecule Inhibitor. Mol Cancer Ther 14, 1837–1847.

Zhang, H., Guttikonda, S., Roberts, L., Uziel, T., Semizarov, D., Elmore, S.W., Leverson, J.D., and Lam, L.T. (2011). Mcl-1 is critical for survival in a subgroup of non-small-cell lung cancer cell lines. Oncogene 30, 1963–1968.

